# Mammalian development does not recapitulate suspected key transformations in the evolution of the mammalian middle ear

**DOI:** 10.1101/023820

**Authors:** Héctor E. Ramírez-Chaves, Stephen W. Wroe, Lynne Selwood, Lyn A. Hinds, Chris Leigh, Daisuke Koyabu, Nikolay Kardjilov, Vera Weisbecker

**Affiliations:** School of Biological Sciences, University of Queensland, Goddard Building 8, St. Lucia 4072, Australia; Division of Zoology, School of Environmental and Rural Sciences, University of New England, NSW 2351, Australia; School of Biosciences, University of Melbourne, VIC 3010, Australia; CSIRO Biosecurity Flagship, GPO Box 1700, Canberra, ACT 2601, Australia; Anatomical Sciences, Adelaide University, North Terrace, SA 5000, Australia; The University Museum, The University of Tokyo, Hongo 7-3-1, Tokyo 113-0033, Japan; Helmholtz-Zentrum Berlin für Materialien und Energie, Hahn-Meitner-Platz 114109 Berlin, Germany

## Abstract

The tympanic ring, malleus and incus of the mammalian middle ear derive from the ancestral primary jaw joint of land vertebrates. In Mesozoic mammals, evolutionary detachment of the mammalian middle ear from the lower jaw occurred when Meckel’s cartilage - the last connection between the middle ear bones and the dentary – disappeared. This disappearance is famously recapitulated in early mammalian development. By extension, it was suggested that other developmental processes also recapitulate specific evolutionary processes that led to mammalian middle ear detachment in Mesozoic mammals. Specifically, developmental posterior/medial displacement and negative allometry of the growing ear ossicles relative to the lower jaw are thought to reflect evolutionary triggers of mammalian middle ear detachment. However, these hypotheses rest on scant developmental data, and have not been tested in a quantitative framework. Here we show, based on μCT scans of developmental series of several marsupials and monotremes, that negative allometry of mammalian middle ear bones relative to the skull occurs only after mammalian middle ear detachment, ruling it out as a developmental detachment trigger. There is also no positional change of ectotympanic or malleus relative to the dentary. The mammalian middle ear bones are differently positioned in two monotreme species, a recent change which is not developmentally recapitulated. Together, our results challenge the developmental prerequisites of previously proposed evolutionary detachment processes, arguing specifically against causal links between mammalian middle ear detachment and brain expansion. Our data are more consistent with a biomechanical detachment trigger relating to the onset of dentary function, but more biomechanical work and palaeontological data are required to test this notion.

## Introduction

The transformation of the middle ear from load-bearing primary jaw joint elements (angular, articular, prearticular, and quadrate) to delicate auditory ossicles (tympanic ring, malleus/gonial, and incus) is one of the oldest and most famous examples of evolutionary developmental biology and represents a textbook example of developmentally informed inference of evolutionary patterns [1, 2]. This transformation – in which several primary jaw joint bones of vertebrates (angular, articular, prearticular, and quadrate) evolved to form the tympanic ring, malleus/gonial, and incus of extant mammals - was famously formulated by 19th century German anatomist B. Reichert [3] based on developmental evidence. Reichert’s work was helped by earlier finds [4] that the anterior leg of the developing mammalian Meckel’s cartilage - which is the cartilaginous precursor of the articular (or body of the malleus [5]) - resides in Meckel’s groove [5], a furrow on the medial side of the dentary (Fig 1 and 2). Meckel’s cartilage constitutes a physical connection between the middle ear and the dentary during mammalian development. The middle ear detaches from the lower jaw when the anterior leg of Meckel’s cartilage is absorbed [6], a process that coincides with a concomitant disappearance of Meckel’s groove.

**Fig 1.**
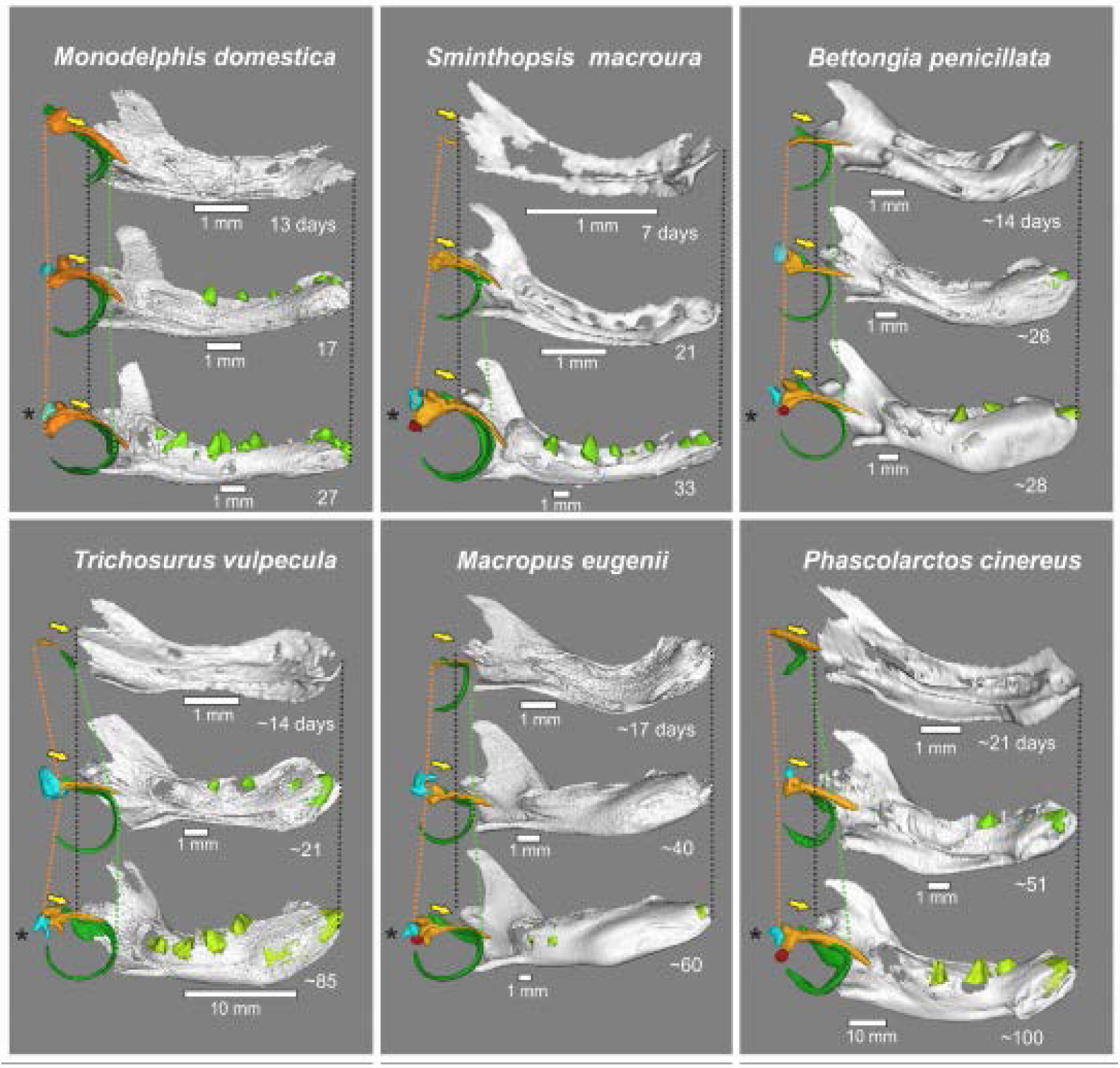
Medial view of middle ear and dentary of selected specimens, each with a young specimen with well-developed Meckel’s groove (top), incipient dental eruption stage (middle), and detached MME (bottom; with asterisk). Orange, malleus; green, ectotympanic; blue, incus; red, stapes; light green, teeth. Lines indicate posteriormost extent of malleus head (orange) and anterior-most rim of the ectotympanic ring (green).

**Fig 2.**
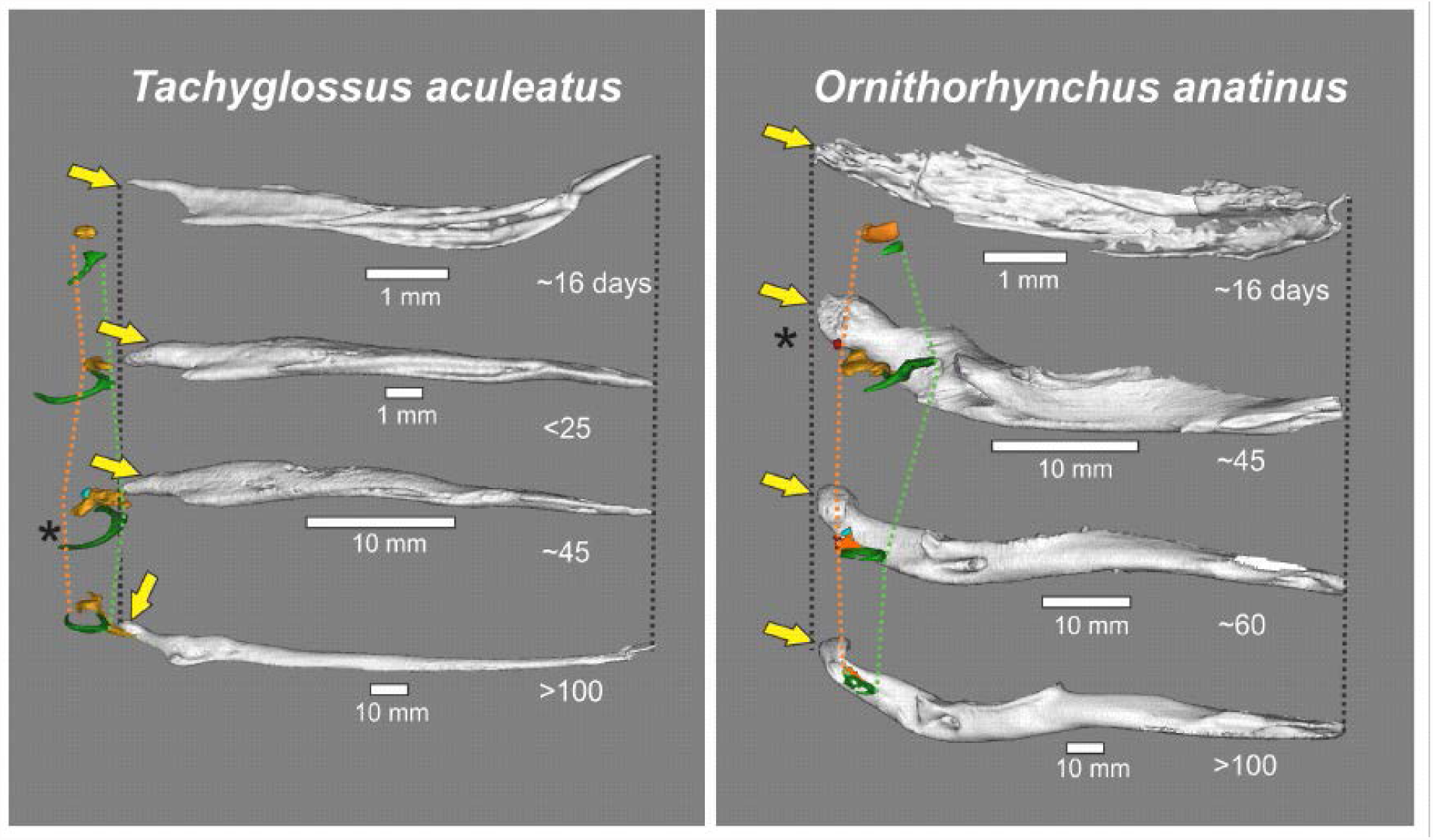
Middle ear and dentary of selected monotreme specimens, from medial view. Orange, malleus; green, ectotympanic; blue, incus; red, stapes; light green, teeth. Lines indicate posterior-most extent of malleus head (orange) and anterior-most rim of ectotympanic ring (green). Asterisk identifies specimens with a detached middle ear and no Meckel’s groove. Note the swing of the *Tachyglossus* dentary from a vertical to a horizontal position.

Disappearance of Meckel’s cartilage appears to be the final step in evolutionary MME detachment just as it is the final step in developmental MME detachment [5]. Due to this, research on the processes behind evolutionary MME detachment is traditionally conducted with extensive reference to developmental observations. For example, lack of Meckel’s groove is considered indicative of MME detachment in fossils [5] as it is in development [6]. Traditional scenarios saw MME detachment as resulting from a conflict between the loadbearing function of the jaw joint and increasing specialization of the postdentary bones for sound conduction, which requires small and gracile bones [7, 8]. This was proposed to result in evolutionary negative allometry of MME bones relative to the jaw and skull, which was thought to be paralleled in mammalian (particularly marsupial) development [5, 9-12]. Based on impressions from mammalian development, particularly older histological reconstructions of developing monotremes [13, 14], it was also suggested that the functional dichotomy between dentary and MME bones resulted in the relocation of the MME bones into a medial position [5, 11, 15].

The notion that MME development might recapitulate homologous evolutionary processes was expanded much further in a study [11] proposing that developmental and evolutionary MME detachment are caused by homologous processes of rapid brain expansion. This “Brain Expansion Hypothesis” (BEH) extensively draws on developmental scenarios in which negative allometry and displacement of the MME relative to the skull are caused by the expanding brain, rather than MME specialization for sound conduction [11]. In Mesozoic mammal evolution, this would have enabled the ossicles to retain suitably small sizes for a sound-conducting role, and as a side-effect triggered the detachment of the MME from the faster-expanding dentary [9-11]. The BEH also predicts that developmental and evolutionary expansion of the mammalian brain displaced the fenestra vestibuli (FV) - through which the ossicular chain connects to the inner ear - into a more posterior position relative to the jaw joint [11]. This was suggested to also cause a displacement of the MME bones posteriorly away from the dentary to maintain the connection between the ossicular chain with the inner ear [11].

The BEH was mostly taken up enthusiastically [5, 16, 17]), despite critical voices from paleontological and developmental studies [18-21]. It is currently an established component of evolutionary scenarios explaining MME detachment [5, 22, 23]. Although middle ear detachment possibly occurred independently in the lineage of monotremes and that leading to the remaining mammals [24, 25], the developmental recapitulation of these processes as specified under the BEH is considered to be essentially the same, with some clade-specific variation in magnitude [5].

Despite broad acceptance of hypotheses matching MME development and evolution, all rest on few quantifiable developmental data, including older histological reconstructions [15, 26]. For example, the only quantitative evidence for developmental negative allometry and displacement of the MME comes from a single developmental series of the opossum, *Monodelphis domestica* [11], whose MME development was controversially [18] correlated with brain growth in the much larger opossum *Didelphis virginiana*. The underlying assumption is that *M. domestica* and *D. virginiana* [17] have the same developmental schedules also gave rise to impressions that MME development occurs at the same time in mammals, roughly 3 weeks after birth [27]. In addition, information on monotremes is also crucial but virtually absent, aside from small-sample histological studies [13, 14, 26].

Here, we provide a large-scale, cross-species investigation of marsupial and monotreme MME development through virtual reconstruction of μCT-scanned developmental series of six marsupial and two monotreme species, focusing on the early-ossifying ectotympanic and malleus (the outer bones of the MME). Marsupials are the organism of choice for studies of therian MME development because of the easy accessibility of young and plesiomorphic arrangement of ME bones, and the notion that newborn marsupials resemble the condition of embryonic mammalian ancestors [7, 12, 27]. By establishing the timing of developmental MME detachment through observations on Meckel’s groove, we characterize the developmental events surrounding MME detachment, and test the developmental basis of claims that evolutionary processes of negative allometry and topological displacement lead to MME detachment in Mesozoic mammals.

## Results

Detachment timing was identified as the age at which Meckel’s groove disappears. This indicates resorption of Meckel’s cartilage and thus detachment of the MME from the middle ear both in MME development and evolution (see introduction). Detachment tends to occur earlier in smaller marsupials (Fig 3). It also tends to occur close to the eruption of molariforms (premolars/molars; Fig 1 and 3). Full ossification of the ossicular chain (including the stapes) is also complete close to the detachment time.

**Fig 3.**
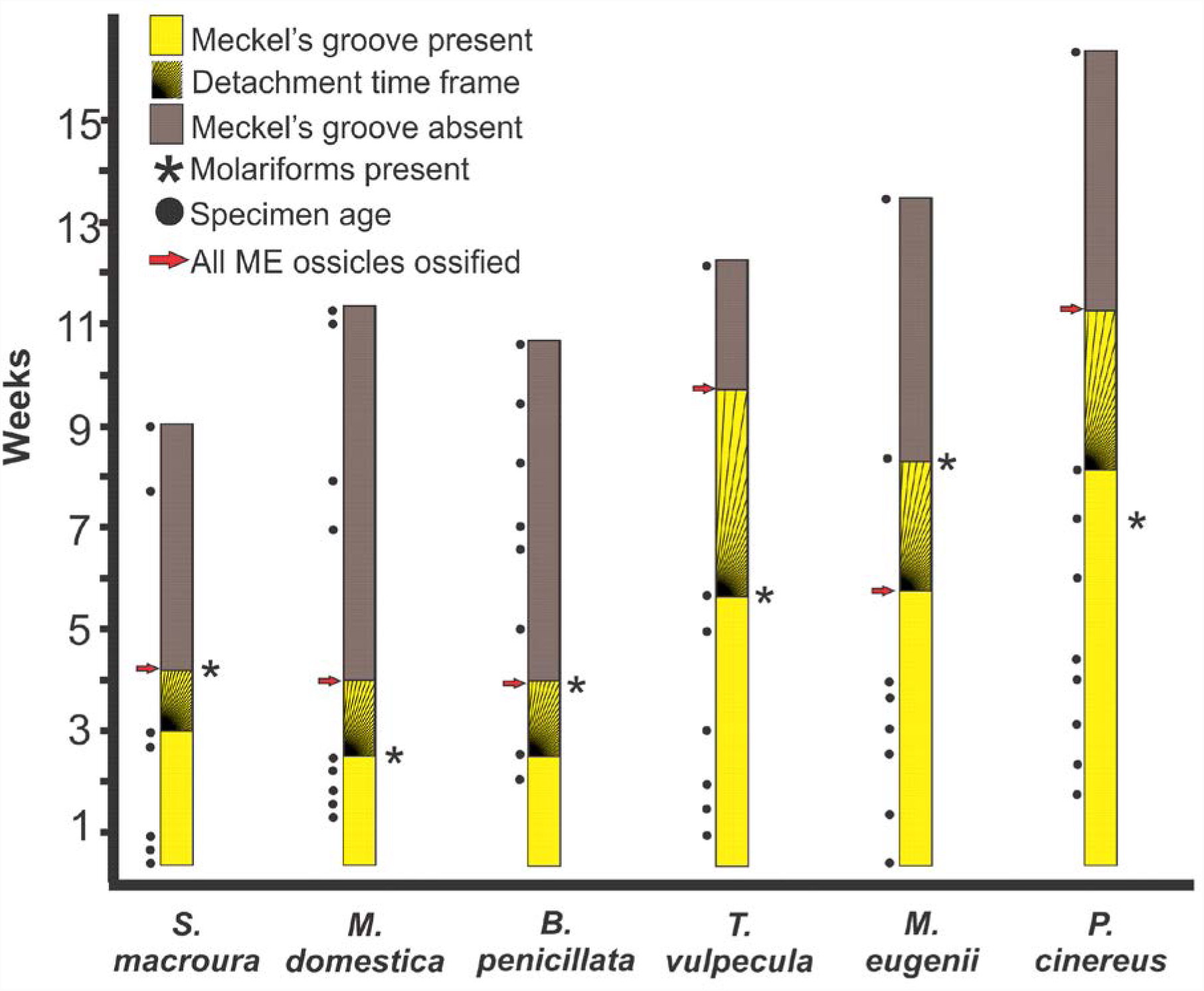
Summary graph of MME detachment timing, as inferred by disappearance of Meckel’s cartilage, in relation to specimen age, dental eruption, and ossicle ossification. Species are ordered from smallest to largest skull length (left to right). “Molariforms” includes premolars and molars. Hatched areas represent gaps between the last specimen with Meckel’s groove and the first specimen without Meckel’s groove, representing the window of time during which MME detachment happens.

Before MME detachment, the ectotympanic and malleus of all species grow with positive allometry relative to both skull and dentary (Fig 4; S1 Table). After MME detachment, the growth of ectotympanic and malleus relative to the skull and dentary is markedly reduced; in the three species with sufficient post-detachment specimens available, a switch to negative allometry (Fig 4; S1 Table) was observed, which seems likely to also occur in the remaining three species sampled (see extrapolations in Fig 4). Both ectotympanic and malleus continue growth until well after MME detachment in all species (Fig 4).

**Fig 4.**
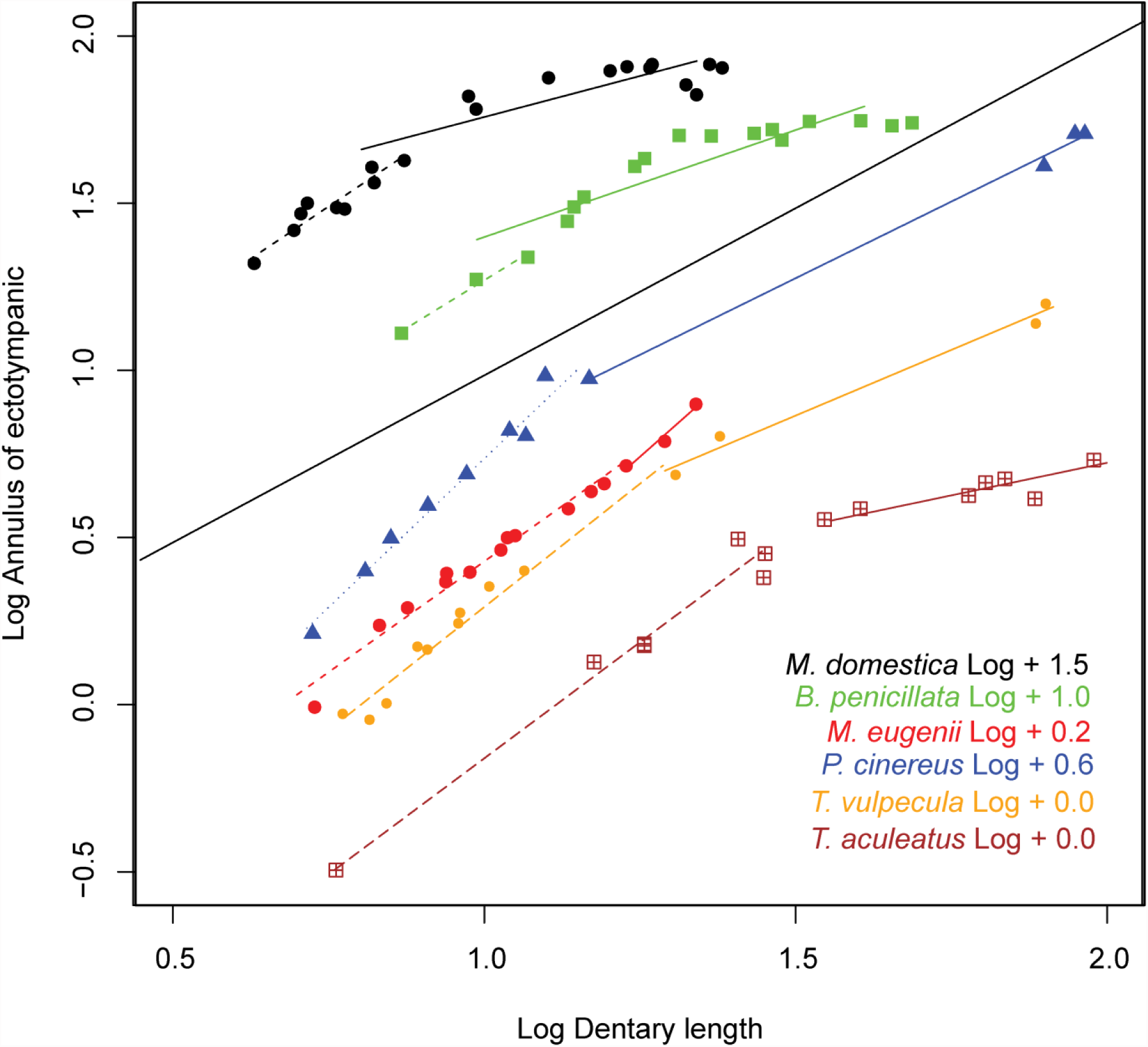
Regressions of ectotympanic annulus dimensions against dentary length in all species sampled, before (dashed lines) and after (solid lines) Meckel’s cartilage detachment. The lines have artificial intercepts added to avoid superimposition. Black line is line of isometry. Post-detachment lines with less than 5 specimens are estimates with the origin of the line forced through the oldest pre-detachment specimens to ensure a conservative estimate of slope.

The positions of ectotympanic or malleus relative to the dentary condyles show little posterior displacement across MME detachment, with slight anterior movements as consequence of ossification more common than slight posterior ones (Fig 1, S2 Table). Medial displacement was also not detected in either monotremes or marsupials, as inter-malleus distances and inter-condylar distances (Fig 1) increase by identical magnitudes (S2 Table) during growth in species of both clades.

Overall, ectotympanic and malleus develop in close association and mostly in their adult position relative to the dentary. This is most evident in the echidna, where dentary, ectotympanic, and malleus together perform a “flipping” movement away from their original vertical position to the adult horizontal orientation (Fig 2). Monotremes are also unusual in that they display divergent positioning of the fenestra vestibuli (FV) relative to the jaw joint: in the platypus, FV and MME ossicles are situated just anterior to the craniomandibular joint, unlike the situation in echidna and most other mammals (Fig 5; S2 Table), where they are positioned well behind the craniomandibular joint.

**Fig 5.**
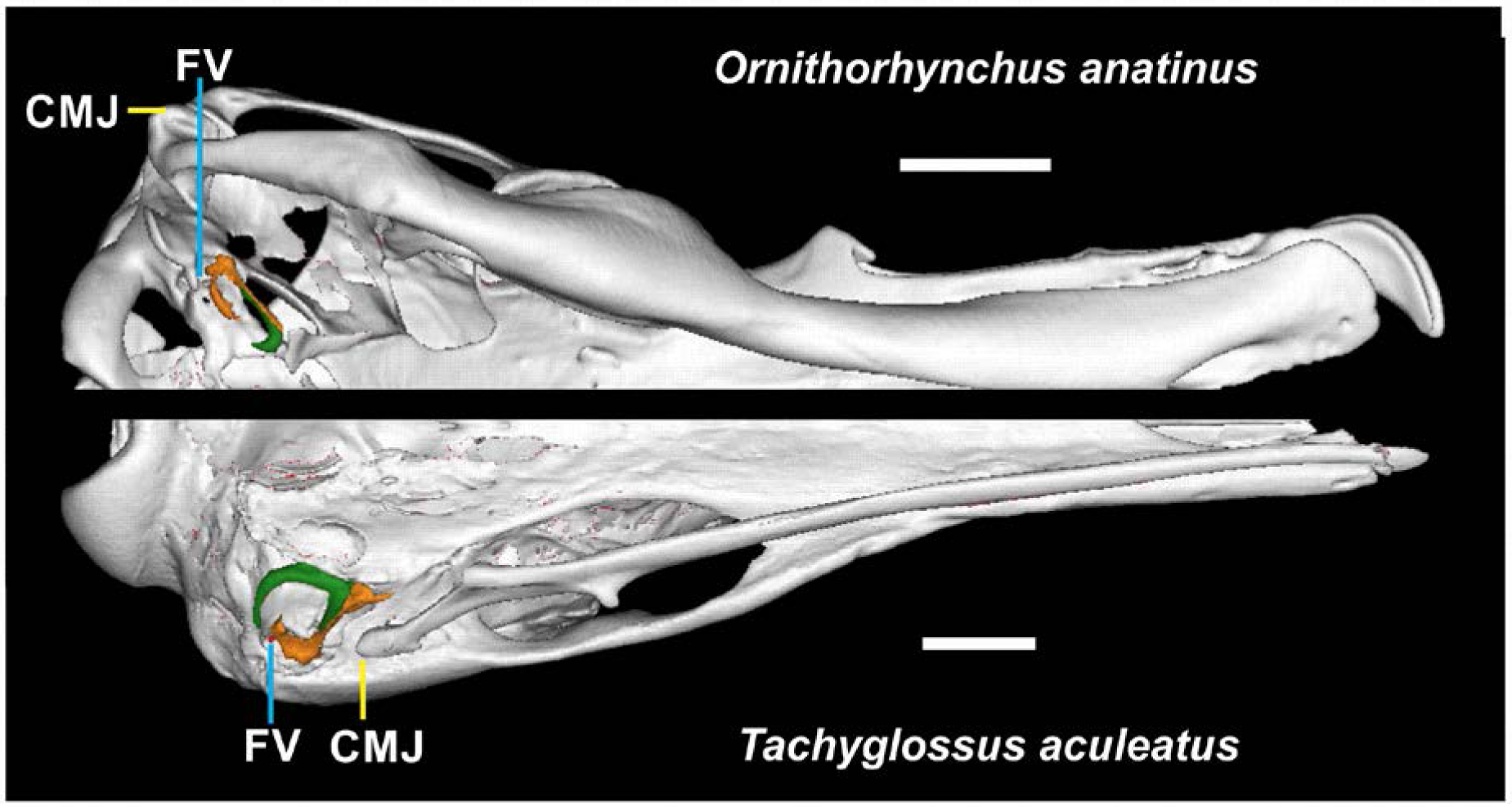
Virtual reconstruction of middle ear region of a platypus (left) and an echidna (right) in ventral view. Note divergent positioning of the fenestra vestibule (FV) and craniomandibular joint (CMJ). Scale bars=10mm.

## Discussion

Our results provide overwhelming evidence against the notion that specific evolutionary processes are recapitulated in MME development. In particular, we found no evidence for hypotheses of detachment-related negative allometry or displacement of the MME bones relative to the skull or jaw joint. Also, contrary to previous suggestions [5], MME development of monotremes and marsupials was overall similar, with exception of the unusual horizontal re-arrangement of the echidna dentary and MME bones. If MME detachment occurred independently in the prototherian and therian lineages [5, 24], this is not obvious through developmental differences. The only between-species differences we found are a clear size-dependence of the timing at which the MME detaches, contrary to suggestions that it occurs at approximately 3 weeks of age in all marsupials [11, 17, 27], and in keeping with the notion that differently-sized mammals have allometric developmental schedules [18].

Negative allometry is the most widely accepted suspected trigger of evolutionary and developmental middle ear detachment in mammals and their ancestors [5, 15, 27]. This is due to the reasoning that the MME bones should have been selected for small sizes that would have improved their ability to conduct sound [11, 25, 28]. However, negative allometry can only be considered a trigger of MME detachment if it occurs before detachment [11]. In contrast, we found negative allometry only after MME detachment in all species investigated. The mechanisms of negative allometry are also not, as previously suggested, related to a stalling of ossicle growth after ossification [27]; both ectotympanic and malleus continue growth until well after MME detachment in all species (Fig 4). Our results contradict the notion that negative allometry is the developmental cause of MME detachment, and an indicator of a similar evolutionary process in Mesozoic mammals [10, 11]. Notably, there is no quantitative information of the relative size of dentary *vs.* post-dentary bones in Mesozoic mammals. A finer-grained investigation of the fossil evidence might be needed to re-assess the temporal relationship of evolutionary scaling patterns in the lower jaw/middle ear region of Mesozoic mammals.

Our results also contradict the assumption [5, 9, 11] that evolutionary topological changes of the MME relative to the jaw joint in a posterior or medial direction are recapitulated in marsupial or monotreme development. Lack of posterior movement in particular contradicts the tenet of the Brain Expansion Hypothesis (BEH) that the developing brain pushes the fenestra vestibuli (FV) – and associated MME bones – into a more posterior position relative to the jaw joint, thus recapitulating an evolutionary process of posterior MME movement [11]. Interestingly, the FV of the platypus is placed slightly anterior to the jaw joint (counter previous suggestions [16]), which is an ancestral condition of earlier synapsids and not normally seen in Mesozoic and extant mammals with a detached middle ear [16]. The existence of an anterior position of the FV in the reasonably large-brained platypus [29] suggests that FV positioning and brain size may not be contingent on each other, and represent separate evolutionary events unrelated to the evolution of the MME. Moreover, FV rearrangement within monotremes must have occurred relatively recently, by earliest in the mid-Miocene, when echidnas and platypus split [30]. It is probably a trait unique to the ancestors of modern platypus, since the Miocene platypus *Obdurodon dicksoni* has the ancestral mammalian condition of the FV well posterior to the jaw joint [31]. Despite this recent transformation, neither species recapitulates a position change of the FV (Fig 2, see also histological evidence in early stages [26, 32]). This casts doubt on the widespread assumption that developmental data can be used to infer topological changes in the much older evolutionary processes of MME detachment in Mesozoic mammals.

In addition to the highly conservative positioning of the MME relative to the jaw joint among our sample, we found that the MME of developing echidnas participates in a gradual re-orientation of the dentary from a vertical to a horizontal position. Such a horizontal arrangement of the MME and dentary is also found in long-beaked echidnas [33]. This adds to the overall impression that the MME and lower jaw represent a tightly integrated developmental unit, rather than two separately developing entities. This matches well with observations on rodents, which suggest that MME and dentary remain connected through collagenous ligaments derived from the remnants of the resorbed Meckel’s cartilage [27, 34]. It is therefore possible that the mammalian middle ear is not be as fully detached from the dentary as currently thought; it may have only exchanged the stiff connection of Meckel’s cartilage with a more flexible ligamentous one.

Our results contradict the widespread notion that MME development can be extrapolated to infer evolutionary MME detachment in Mesozoic mammals. They specifically challenge the developmental premises of the “Brain Expansion Hypothesis”, supporting fossil-based suggestions [20] that MME detachment and brain expansion represent separate events at a time when mammalian characteristics rapidly arose [35]. In contrast, the consistent timing of MME detachment with molariform eruption and full MME ossification suggests that developmental MME detachment relates to a switch in function, where the MME assumes its sound-conducting role and the dentary takes over biomechanical loading related to mastication [25, 28], or sucking [15]. This broadly supports more traditional functional explanations of MME evolution, which also focused on changes in dentary function as triggers of MME detachment [15, 25, 36]. However, the unexpectedly narrow limits of developmental inference found here caution against any interpretation of development in the absence of quantitative palaeontological data. This is a challenge for the field because fossils are scarce and scattered across the large phylogeny of Mesozoic mammals. Widespread convergence of MME anatomy in Mesozoic mammals [20] also hampers quantitative interpretations of evolutionary MME detachment. However, all hypotheses of MME detachment rest on some form of mechanical driver (either brain expansion or mastication), so that it might be possible to infer more detail on MME functionality using biomechanical modelling. Further fossil evidence, as well as more detailed work on the biomechanics of the developing and evolving jaw joint, may therefore improve resolution on this issue.

## Materials and Methods

The sample contained two monotremes (the platypus, *Ornithorhynchus anatinus* and the echidna, *Tachyglossus aculeatus*) and six marsupial species, including one didelphimorphian (the gray short-tailed opossum *Monodelphis domestica*), one dasyuromorphian (the stripe-faced dunnart *Sminthopsis macroura*), four diprotodontians (the woylie *Bettongia penicillata*, the tammar wallaby *Macropus eugenii*, the koala *Phascolarctos cinereus*, and the brush-tailed possum *Trichosurus vulpecula*; for accession numbers and age estimates, see S3 Table). Most marsupial specimens were fixed and stored in buffered formalin, except for the earlier stages of *M. domestica* (postnatal day 10-17) which were clear-stained specimens and later stages of *M. domestica* (day 27 onwards) which were frozen and/or ethanol-preserved [37]. Monotremes were obtained from ethanol-preserved museum specimens. Specimens were scanned using micro-computed tomography (μCT) scanners at the Helmholtz-Zentrum in Berlin, the Cambridge University Department of Engineering, University College London, and the Centre for Advanced Imaging at the University of Queensland. Scanning involves the transmission of X-rays through the specimens under different rotation angles; in this way 2D angular projections are collected over 180° or 360°. The set of projections is used for 3D reconstruction of the matrix of absorption coefficients in the sample by a back-projection algorithm. The achieved spatial resolution is in the range of dozens to a few hundred micrometers.

The time of detachment of the MME bones from the dentary was documented through determining the presence or absence of Meckel’s groove, which is the last point of contact between the dentary and the middle ear (see text). Ossification status of the auditory chain and eruption of molariforms was also documented to provide a timeframe reference for MME detachment relative to other developmental events.

## Allometry analyses

To test the hypothesis that MME detachment might be due to negative allometry of the MME bones relative to the remainder of the skull, we measured proxies of cranial size (condylobasal length, dentary length) and proxies of middle ear size (bullar part of ectotympanic diameter (only available in marsupials), the diameter of the ectotympanic ring, and the anteroposterior length of the gonioarticular portion of the malleus (S1 Fig; S1 Table and S4 Table). Allometric coefficients between ectotympanic and malleus lengths against condylobasal and dentary length were determined using Standardized Major Axis (SMA) regression in the software package R [38] and the package ‘smatr’ [39]. Negative or positive allometry was identified when the coefficients were statistically significantly higher or lower than unity, respectively, with α = 0.01 [40]. Pre-detachment analyses were not performed for *O. anatinus* and *B. penicillata*, due small sample size. Only *B. penicillata* and *T. aculeatus* had sufficient specimen representation for a quantification of post-detachment allometry. We therefore only visually evaluated the position of data points relative to the pre-detachment slope in the other species, and estimated the slope of post-detachment data regressions that were forced through the oldest pre-detachment specimen. We also included assessments of allometry in the external auditory meatus of marsupials, although specimen numbers limit the reliability of this regression as it is formed well after detachment of middle ear bones in most of the species evaluated.

## Tests for ontogenetic middle ear displacement from the dentary

To assess a possible anteroposterior separation of the MMME from the dentary-squamosal (D-SQ) jaw joint, we measured the distance between the anterior-most part of the dentary up to the anterior part of the ectotympanic (ECT), the distance between the anterior-most part of the dentary and the posterior part of the malleus (MAL), and the distance of the anteroposterior length of the dentary at the level of the condyloid process (CON) (S1 Fig). We assigned to CON a value of one and transformed ECT and MAL as a proportion of CON. ECT/MAL values greater and smaller than 1 indicate movements to a more anterior and posterior position, respectively, relative to the dentary condyle (S2 Table).

The suggestion of developmental mediolateral displacement of MME bones from the dentary was tested by regressing the distances between the heads of the two mallei against the distance between the condyloid processes of each dentary (measured as in S1 Fig). For each species, these measurements were compared using Spearman’s rank correlations (S2 Table). We expected a negative correlation in case of medially-directed movement of the malleus relative to the dentary.

## Acknowledgments

We thank C. Kemper and D. Stemmer (Mammal Collection of the South Australian Museum) for access to *Bettongia* and monotreme specimens, and H. Janetzki (Queensland Museum), D. Pickering (Museum Victoria), and S. Ingleby (Australian Museum) for access to monotreme specimens. We thank K. Mardon (Centre for Advanced Imaging, University of Queensland) for help with μCT scanning. Many thanks to T. Rowe and T. Macrini for providing post-detachment *Monodelphis* specimens.

## Supporting information captions

**S1 Fig. Measurements taken for the analyses.** 1, Malleus length; 2, tympanic annulus diameter; 3 diameter of bullar part of ectotympanic; 4, skull length; 5, dentary length; 6, inter-malleus length; 7, intercondylar length; 8, Distance from tip to dentary to anterior rim of ectotympanic; 9, distance from dentary tip to posterior end of malleus head.

**S1 Table. Summary of log-log bivariate regressions for four species of marsupials and one monotreme, before and after detachment of middle ear bones, regressing dentary length and condylobasal (CB) lengths against dimensions of the bullar part of the ectotympanic (bullar part), annulus of the ectotympanic (annulus), antero-posterior length of the malleus (malleus a.-p length), and external auditory meatus (EAM).** N, Sample size; R2, adjusted coefficient of correlation; slope, coefficient of allometry under Standardised Major Axis (SMA); P, P-value for SMA regressions; P1, probability that the slope is equal to 1. Significant allometric slopes are in bold.

**S2 Table. Tests for posterior and medial displacement of the post-dentary bones during development.** Antero-posterior positioning defines the dentary length (distance between dentary tip and dentary condyle) as one and determining the distances between dentary tip and ectotympanic/malleus as a percentage of dentary length. Light grey fields identify overall posterior movement of ossicles relative to dentary condyle, dark grey fields identify overall anterior movement; white fields suggest no change.

**S3 Table. Accession number, age, references for aging of the specimens used in this study.** *Bettongia penicillata*; South Australian Museum (SAM). *Macropus eugenii*; CSIRO Sustainable Ecosystems Canberra, colony collected (ACT permit K1606); CSIRO Sustainable Ecosystems Animal Ethics Approval 06-36. *Phascolarctos cinereus*; Adelaide University, field collected (DENWR (SA) License Nos. K23749 /1 to 25). *Sminthopsis macroura*, collected from a colony (MAEC (Vic) License No. 06117). *Trichosurus vulpecula*, collected from a colony (MAEC (Vic) License No. 06118). *Monodelphis domestica*, collected from preserved specimens University Museum of University of Tokyo (UMUT), and scans courtesy of by T. Rowe and T. Macrini. *Ornithorhynchus anatinus*, collected from preserved specimens at Queensland Museum (J and JM), Australian Museum (AusMusM). *Tachyglossus aculeatus*, collected from preserved specimens at Queensland Museum (J and JM), Australian Museum (AusMusM), and South Australian Museum (SAM).

**S4 Table. Measurements used for the analyses. Maximum distance between the posterior part of the malleus (PPM).** Maximum distance of the condyloid process (CDP). Anterior-most part of the dentary up to the anterior part of the ectotympanic (ECT). Anterior-most part of the dentary and the posterior part of the malleus (MAL). Anteroposterior length of the dentary at the level of the condyloid process (CON). Dentary length (DL). Condylobasal length (CBL). Malleus antero-posterior length (MP). Bullar part of ectotympanic (BPE). Annulus of ectotympanic (ANE). Auditory meatus (AM).

